# Regenerative capacity in the lamprey spinal cord is not altered after a repeated transection

**DOI:** 10.1101/410530

**Authors:** Kendra L. Hanslik, Scott R. Allen, Tessa L. Harkenrider, Stephanie M. Fogerson, Eduardo Guadarrama, Jennifer R. Morgan

## Abstract

The resilience of regeneration in vertebrate tissues is not well understood. Yet understanding how well tissues can regenerate after repeated insults, and identifying any limitations, is an important step towards elucidating the underlying mechanisms of tissue plasticity. This is particularly challenging in tissues such as the nervous system, which contain a large number of terminally differentiated cells (i.e. neurons) and that often exhibits limited regenerative potential in the first place. However, unlike mammals that exhibit very little spinal cord regeneration, many non-mammalian vertebrate species, including lampreys, fishes, amphibians and reptiles, regenerate their spinal cords and functionally recover even after a complete spinal cord transection. It is well established that lampreys undergo full functional recovery of swimming behaviors after a single spinal cord transection, which is accompanied by tissue repair at the lesion as well as axon and synapse regeneration. Here, using the lamprey model, we begin to explore resilience of spinal cord regeneration after a second spinal re-transection. We report that by all functional and anatomical measures tested, the lampreys regenerated after spinal re-transection just as robustly as after single transections. Recovery of swimming behaviors, axon regeneration, synapse and cytoskeletal distributions, and neuronal survival were nearly identical after a single spinal transection or a repeated transection. Thus, regenerative potential in the lamprey spinal cord is largely unaffected by spinal re-transection, indicating a greater persistent regenerative potential than exists in some other highly-regenerative models. These findings establish a new path for uncovering pro-regenerative targets that could be deployed in non-regenerative conditions.

## Introduction

High regenerative capacity has demonstrated in a number of invertebrate and vertebrate animals. Classic models for whole body regeneration include hydras, planarians, and many annelids, which can regenerate entire animals with proper body form from tiny pieces of tissues or after repeated amputations (Reddien and Sanchez Alvarado, 2004; Tanaka and Reddien, 2011; Bely, 2014). Similarly, many instances of organ and tissue regeneration have been observed amongst vertebrate species. For example, zebrafish can regenerate complex tissues and organs including the heart, liver and fins (Gemberling et al., 2013; Wang et al., 2017). Mexican axolotl salamanders are known to regenerate their limbs, tails, skin, and several internal organs (Muneoka and Bryant, 1982; Echeverri and Tanaka, 2002; Monaghan et al., 2007; Voss et al., 2009; McCusker and Gardiner, 2011; Seifert et al., 2012; Voss et al., 2013; Erickson et al., 2016). Other amphibians such as *Xenopus* tadpoles can regenerate spinal cord, limb buds, tail and lens (Slack et al., 2008; Gibbs et al., 2011). This regenerative capacity is not limited to non-mammalian vertebrates, as neonatal mice can regenerate digit tips and heart (Reginelli et al., 1995; Uygur and Lee, 2016; Dolan et al., 2018).

Remarkably, even tissues with a large number of terminally differentiated cells, such as the central nervous system, can readily regenerate in vertebrates. As examples, zebrafish and amphibians can regenerate parts of their retina, optic nerve, and brain (Sperry, 1947; Fawcett and Gaze, 1981; Vergara and Del Rio-Tsonis, 2009; Goldshmit et al., 2012; Gorsuch and Hyde, 2014; Morgan and Shifman, 2014; Williams et al., 2015). Species ranging from lampreys and fishes to amphibians and reptiles can regenerate spinal cord structures (Tanaka and Ferretti, 2009; Zukor et al., 2011; Goldshmit et al., 2012; Diaz Quiroz and Echeverri, 2013; Bloom, 2014; Morgan and Shifman, 2014). Although central nervous system (i.e. brain and spinal cord) regeneration is poor in mammals, peripheral nerve regeneration is particularly robust in most vertebrates, including adult mammals (David and Aguayo, 1981; Son and Thompson, 1995; Kang and Lichtman, 2013). While these and many other examples of successful regeneration have been demonstrated across the animal kingdom for over a century, what is not understood is how well regenerative capacity persists after repeated injuries.

Repeated rounds of injury and recovery have been followed in only a small number of experimental models, with surprisingly varied outcomes on regenerative capacity. At one extreme, whole planarians can regenerate entire body structures from as little as 1/279^th^ of the original parent animal (Morgan, 1898; Newmark and Sanchez Alvarado, 2002). Because planarians reproduce by fission, they can survive repeated rounds of resection and regeneration and are therefore are technically immortal. Likewise, the zebrafish caudal fin can undergo repeated cycles of normal regeneration even after 27 amputations at the same location (Azevedo et al., 2011). At the other extreme, salamanders and newts exhibit imperfect regeneration of limb structures beginning with the second amputation (Dearlove and Dresden, 1976; Frobisch et al., 2014; Bryant et al., 2017). Therefore, regenerative capacity is limited in certain cases.

In comparison to the examples described above, very little if anything is known about how regeneration of nervous system tissues is affected by repeated injuries. Yet, understanding the extent of regenerative capacity in the spinal cord or brain could provide important insights into the mechanisms of nervous system plasticity, as well as the limitations that occur in higher vertebrates such as mammals. To begin testing the resilience of regenerative capacity in the vertebrate nervous system, we followed the behavioral and anatomical outcomes after two successive spinal cord transections in sea lampreys, *Petromyzon marinus*. Lampreys undergo full functional recovery of swimming behaviors within 10-12 weeks after completely transecting, or severing, the spinal cord (Rovainen, 1976; Selzer, 1978; Cohen et al., 1986; Davis et al., 1993; Oliphint et al., 2010). Behavioral recovery is accompanied by tissue repair at the lesion site, regeneration of descending and ascending axons several millimeters beyond the lesion site, and formation of new synaptic connections (Rovainen, 1976; Wood and Cohen, 1981; Yin and Selzer, 1983; Mackler and Selzer, 1985; Davis and McClellan, 1994; Oliphint et al., 2010). Amongst the descending neurons are 32 identified “giant” reticulospinal neurons, which reside in stereotypical locations in the lamprey midbrain and hindbrain, with known probabilities of survival and regeneration. Some are reproducibly “good survivors/regenerators,” while others are “poor survivors/regenerators”, a unique feature of the lamprey model that allows for determination of regenerative capacity at the level of individual neurons (Jacobs et al., 1997; Shifman et al., 2008; Busch and Morgan, 2012; Lau et al., 2013; Barreiro-Iglesias, 2015). In this study, we measured functional and anatomical recovery after an initial spinal cord transection and also after a second spinal re-transection at the same lesion site. We report here nearly identical behavioral recovery, tissue repair, and neural regeneration after spinal transection and re-transection, indicating that spinal cord regeneration in lampreys is resilient to repeated injuries.

## Materials and methods

### Spinal cord surgeries

Spinal cord transections were performed as previously described (Oliphint et al., 2010; Lau et al., 2011; Fogerson et al., 2016). Briefly, late stage larval sea lampreys (*Petromyzon marinus*; 11-13 cm) were first anesthetized in 0.2 g/L MS-222 (Tricaine-S; Western Chemical, Inc.; Ferndale, WA). Once anesthetized, a small horizontal incision was made at the level of the 5^th^ gill through the skin and muscle to reveal the spinal cord, after which it was completely transected using fine iridectomy scissors. The spinal transection was visually confirmed, and the incision was subsequently closed with sutures (Ethilon 697G Ethilon Nylon Suture; Ethicon US, LLC; Somerville, NJ). Animals were housed post-operatively in isolated tank breeders within 10-gallon aquaria and held at room temperature (RT; 20-25°C). At 11 weeks post-injury (wpi), the regenerated lampreys were re-anesthetized, and their spinal cords were re-transected through the original lesion scar using the same procedure. After spinal re-transection, the lampreys were allowed to recover for another 11 wpi prior to tissue harvest. All procedures were approved by the Institutional Animal Care and Use Committee at the Marine Biological Laboratory in accordance with standards set by the National Institutes of Health.

### Behavioral analysis

After spinal transection or re-transection, the lampreys’ swimming movements were scored twice per week during the recovery periods, as previously described (Oliphint et al., 2010; Herman et al., 2018). The scoring criteria were as follows: 0 – immediately post-operatively, lampreys exhibited no response to a light tail pinch; 1 – only head movements were observed; no tail movements occurred below the lesion site; 2 – brief periods of self-initiated swimming occurred, but with atypical movements and body shapes; 3 – lampreys demonstrated longer periods of swimming with more normal undulations and fewer abnormalities; 4 – lampreys exhibited persistent bouts of swimming with normal sinusoidal undulations that were comparable to uninjured, control lampreys. The average movement scores were calculated for n=10-18 animals and graphed in GraphPad Prism 6.0c (GraphPad Software, Inc.; La Jolla, CA). Additionally, at 1, 3, and 11 wpi, during both recovery periods, still images of the lampreys were acquired using a Sony Handycam HDR-CX455.

### Spinal cord dissection and bright field imaging

At the appropriate post-injury time points, lampreys were re-anesthetized, and ∼4 cm segments of the spinal cords surrounding the lesion site were microdissected in fresh, oxygenated lamprey Ringer: 100 mM NaCl, 2.1 mM KCl, 1.8 mM MgCl_2_, 4 mM glucose, 2 mM HEPES, 0.5 mM L-glutamine, 2.6 mM CaCl_2_, pH 7.4. For most experiments, the spinal cords were fixed immediately in 4% paraformaldehyde (PFA) in 0.1 M phosphate buffered saline (PBS, pH 7.4) for 3 hours at RT and then overnight at 4°C, followed by washing for 3 x 5 min with 0.1 M PBS (pH 7.4). Brightfield images of spinal cords at 1, 3, and 11 wpi were acquired at 30X magnification using a Zeiss Axiocam503 color camera mounted to a Zeiss Axio Zoom.V16 fluorescence stereo zoom microscope (1X, 0.25 NA Zeiss Plan-NeoFluar Z objective).

### Anterograde labeling of regenerated axons

For a subset of experiments, bulk anterograde labeling of regenerated axons was performed in transected (n=14) and re-transected (n=7) spinal cords, as previously described (Lau et al., 2013). Axons were labeled with a fluorescent dye (5 mM Alexa Fluor® 488-conjugated dextran; 10 kDa; Thermo Fisher, Inc.), diluted in lamprey internal solution (180 mM KCl, 10 mM HEPES, pH 7.4) via a 1×1×1 mm piece of Gelfoam (Pfizer; New York, NY), which was applied 5 mm rostral to the lesion site. After application, the dye was allowed to transport for 3-6 days before harvesting the spinal cords. Labeled spinal cords were imaged live in lamprey Ringer. Confocal Z-stacks were collected using a Zeiss LSM 510 laser scanning confocal attached to an Axioskop 2FS upright microscope (10X, 0.3 NA Zeiss Plan-NEOFLUAR objective). Maximum intensity projections of the spinal cords, ranging from ∼1 mm proximal to 5 mm distal to the lesion, were generated using the Zeiss LSM software. After stitching the projections together in Adobe Photoshop, the number of labeled, regenerated axons was counted at 1 mm intervals starting from the center of the lesion. Graphing and statistical analyses were conducted using GraphPad Prism 6.0c (GraphPad Software; La Jolla, CA).

### Immunofluorescence assays

For immunofluorescence experiments, fixed spinal cords were cryoprotected in 12%, 15%, and 18% sucrose in 0.1 M PBS, pH 7.4 for ≥ 3 hours each at RT, or overnight at 4°C. A 2-cm length of each spinal cord was then embedded horizontally in O.C.T. Compound (EM Sciences; Hatfield, PA). Longitudinal sections (14 μm thickness) were collected onto Superfrost Plus microscope slides (Fisher Scientific; Pittsburgh, PA) using a Leica CM1850 cryostat and stored at −20°C until use.

Cryosections were incubated in blocking buffer containing 9.5% normal goat serum (Life Technologies, Carlsbad, CA) and 0.5% Triton-X 100 for 45 minutes at RT. Next, the sections were incubated in primary antibodies diluted 1:100 in an antibody signal enhancer (ASE) solution for 2 hours at RT, as described in Rosas-Arellano et al., 2016. The primary antibodies used for this study were: a mouse monoclonal antibody raised against lamprey neurofilament-180 (LCM16; kind gift from Dr. Michael Selzer); a mouse monoclonal SV2 antibody that was deposited to the DSHB by Dr. Kathleen Buckley (DSHB; Iowa City, IA) (Buckley and Kelly, 1985); and a mouse monoclonal anti-α-Tubulin antibody (clone DM1A; Sigma-Aldrich; St. Louis, MO). The NF-180 antibody (Jin et al., 2009) and SV2 antibody (Bloom et al., 2003; Oliphint et al., 2010; Lau et al., 2011; Busch and Morgan, 2012; Busch et al., 2014) have been extensively characterized in lamprey nervous tissues. The α-tubulin antibody is further characterized in Supplemental Fig. 1. After primary antibody incubation, the sections were washed for 3 x 10 minutes at RT in wash buffer (20 mM Na phosphate buffer pH 7.4, 0.3% Triton X-100, 450 mM NaCl), followed by a 1 hour incubation at RT in secondary antibody (1:300 Alexa Fluor™ 488-conjugated goat anti-mouse IgG (H+L); ThermoFisher Scientific). For labeling of actin cytoskeleton, sections were directly labeled with Acti-Stain™ 488 phalloidin (Cytoskeleton, Inc.; Denver, CO) at 1:200 diluted in blocking buffer for 45 min at RT. Finally, sections were washed in wash buffer for 3 x 5 min, followed by 15 min in 5 mM Na phosphate buffer, pH 7.4. Slides were then mounted with ProLong® Gold antifade reagent with DAPI (Life Technologies, Inc.) in order to label nuclei. After immunostaining, sections were imaged in ZEN 2.3 software using a Zeiss Axiocam 503 color camera mounted onto a Zeiss Axio Imager.M2 upright microscope (10X, 0.3 NA and 40X, 1.3 NA Zeiss EC Plan-Neofluar objectives).

**Figure 1.**
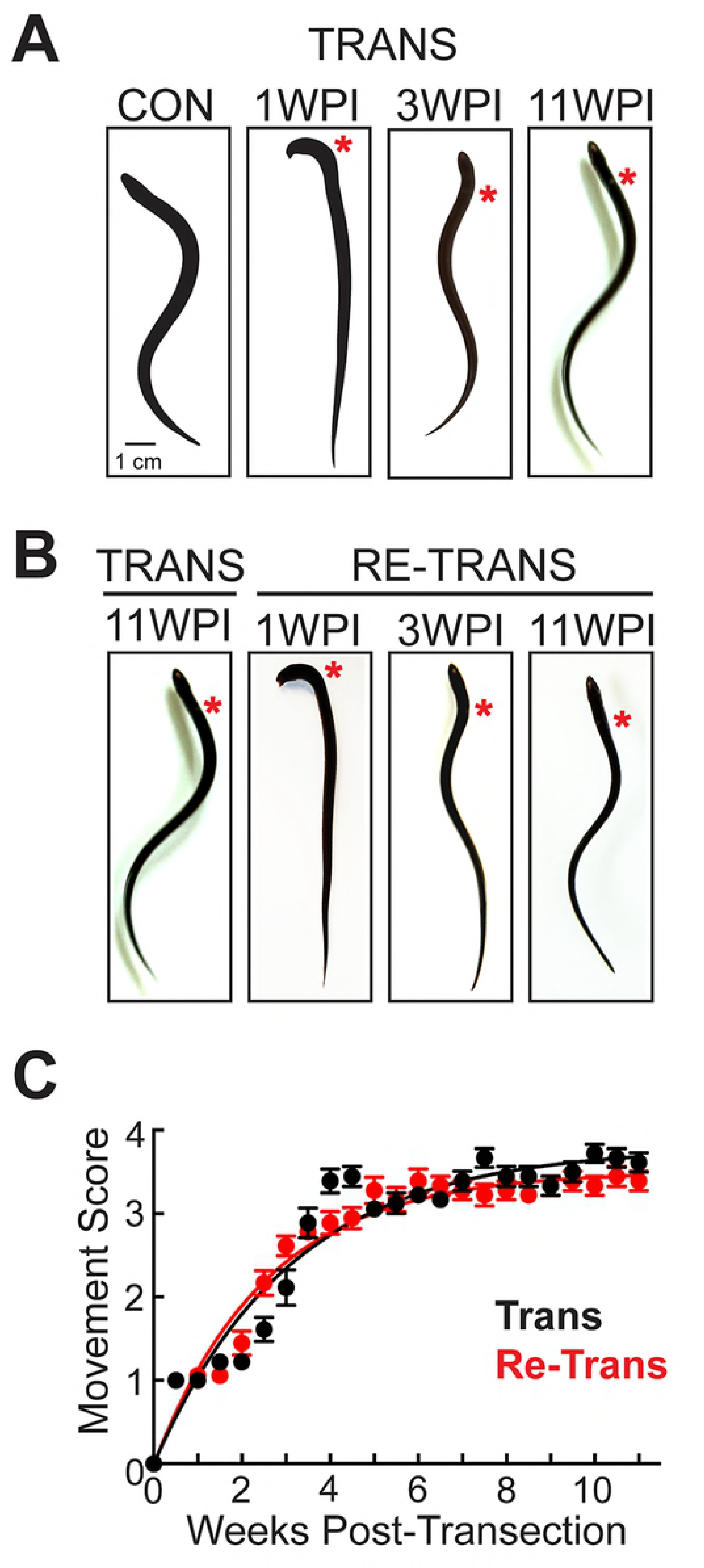
Functional recovery of swimming in lampreys after two successive spinal cord transections. **(A-B)** Still images of lampreys illustrating several distinct stages of functional recovery after spinal transection (A) or re-transection (B). The body shapes are similar at each post-injury time point. Scale bar in panel A also applies to B. **(C)** Time course of functional recovery of swimming movements in transected versus re-transected lampreys shows no obvious difference. Data points represent mean ± SEM from n=18 lampreys. Data were well fit by an exponential curve (Trans R^2^=0.78; Re-Trans R^2^=0.78).

### Retrograde labeling of regenerated neurons

Regenerated giant RS neurons were retrogradely labeled, as previously described (Shifman et al., 2008). Briefly, at 11 wpi, the regenerated axons in the transected and re-transected spinal cords were labeled by inserting a 1×1×1 mm pledget of Gelfoam soaked in 10 mM tetramethylrhodamine dextran (TMR-DA; 10 kDa; ThermoFisher) into the spinal cord at a location 5 mm caudal to the lesion site (Shifman, et al., 2008). The TMR-DA was allowed to retrogradely transport for 9 days prior to harvesting the brains. Brains were imaged live using a Zeiss Axio Zoom.V16 fluorescence stereo zoom microscope. Giant RS neurons that had regenerated their axons distal to the lesion were identified by their fluorescently-labeled cell bodies, while giant RS neurons that did not regenerate were devoid of dye. Mean fluorescence intensity for each RS neuron was measured using ImageJ software. The percentage of regenerated RS neurons was determined from n=10 lampreys for each experimental condition. All graphing and statistical analyses were performed in OriginPro 7.0 (OriginLab Corp.; Wellesley, MA).

### Nissl staining

After imaging the regenerated neurons, the lamprey brains were subsequently fixed overnight at 4°C in 4% PFA in 0.1M PBS, pH 7.4, washed 3 x 15 min with 0.1M PBS (pH 7.4) and stained with Toluidine Blue O (EM Sciences; Hatfield, PA) to label the Nissl substance (Shifman et al., 2008; Busch and Morgan, 2012). Brains were incubated in 1% Toluidine Blue O solution containing 1% borax (pH 7.6) for 20 minutes at 37°C. The brains were then destained in Bodian’s fixative (72% EtOH; 5% glacial acetic acid; 5% formalin) until the desired tissue contrast was obtained. Next, the brains were dehydrated in 95% and 100% ethanol (2 x 5 min each) and cleared in cedarwood oil at 65°C for 2 hours prior to mounting on slides with Permount. Brightfield images of whole brains and giant RS neurons were acquired at 20-80x magnification using a Zeiss Axio Zoom.V16 fluorescence stereo zoom microscope.

Image analysis was performed using ImageJ software. The mean intensity of each giant neuron was measured. Cells with a positive mean intensity after background subtraction were labeled Nissl (+), and cells with a negative mean intensity were labeled Nissl (-). The average percentage of Nissl (+) cells was plotted by cell type and for the overall brain from n=9 transected, n=10 re-transected lampreys. Graphing and statistical analyses were performed in OriginPro 7.0 (OriginLab Corp.; Wellesley, MA).

## Results

### Lampreys exhibit robust functional recovery after spinal re-transection

The goal of this study was to determine the extent to which lampreys can functionally recover and regenerate their spinal cord structures after a repeated transection. We thus began by observing their behavioral recovery after two successive spinal transections. First, the lampreys were spinally transected at the level of the 5^th^ gill, after which they were allowed to recover for 11 weeks post-injury (wpi). At 1 wpi, the lampreys were paralyzed below the lesion site, and only head movements were observed (Fig. 1A). At 3 wpi, the lampreys regained their ability to swim but displayed abnormal movements such as rapid head oscillations, abnormal body contractions, and shallow sinusoidal waves (Fig. 1A). Once the lampreys reached 11 wpi, they exhibited normal sinusoidal swimming movements (Fig. 1A). After this initial recovery period, lampreys underwent a second spinal transection at the original lesion site and were subsequently allowed to recover for another 11 wpi. Re-transected lampreys recovered along the same timeline and displayed similar locomotor behaviors (Fig. 1B). During both recovery periods, the swimming behaviors were recorded twice per week using an observational movement scoring, as described in Oliphint et al., 2010, where a score of 0 indicates complete paralysis; 1 indicates head wagging, but no forward movement; 2 indicates brief bouts of abnormal, self-initiated swimming; 3 indicates longer durations of persistent swimming with more regular movements; 4 represents normal sinusoidal swimming. The movement scoring indicated that both transected and re-transected lampreys recovered robustly along similar trajectories. Singly transected lampreys recovered to ∼90% of normal levels by 11 wpi (Fig. 1C). The recovery curve was well fit by an exponential process that reached a half maximum (t_1/2_) at 2.6 ± 0.2 wpi (R^2^ = 0.78, n=18 lampreys), which was similar to previous reports (Fig. 1C) (Oliphint et al., 2010; Herman et al., 2018). After spinal re-transection, this same cohort of lampreys recovered to ∼85% of normal swimming movements by 11 wpi, reaching half maximum (t_1/2_) at 2.4 ± 0.1 wpi (R^2^ = 0.78, n=18 lampreys) (Fig. 1C). Thus, remarkably, lampreys were able to recover normal swimming movements to the same degree after multiple spinal transections.

### Lesion repair is complete after spinal re-transection

Next, we examined tissue repair in the lamprey spinal cord after transection and re-transection. Uninjured, control spinal cords are translucent and well organized with several giant axons (gray lines) and large neurons (gray spheres) visible along the longitudinal axis (Fig. 2A). After the initial spinal transection, at 1 wpi, the proximal and distal stumps of the spinal cord were largely disconnected, joined only by a thin layer of meninges, and the central canal (red arrow) was swollen (Fig. 2B). At 3 wpi, the proximal and distal stumps had become re-connected by a glial-ependymal scar (Fig. 2C) (Selzer, 1978; Lurie et al., 1994). The central canal could also be seen extending through the lesion, still swollen. By 11 wpi, the tissue regained a more normal translucent appearance, and the lesion scar appeared more healed, though the central canal remained swollen (Fig. 2D-E). After spinal re-transection, the spinal cords exhibited similar morphologies at these post-injury time points (Fig. 2E-H). A notable exception was at 1 wpi, where the re-transected spinal cords routinely exhibited an advanced development of tissue repair at the lesion site (Fig. 2F). The re-transected spinal cords were also slightly narrower at the lesion site (Fig. 2F-H). Thus, in addition to behavioral recovery, there was robust repair of the spinal cord tissues in re-transected lampreys.

**Figure 2.**
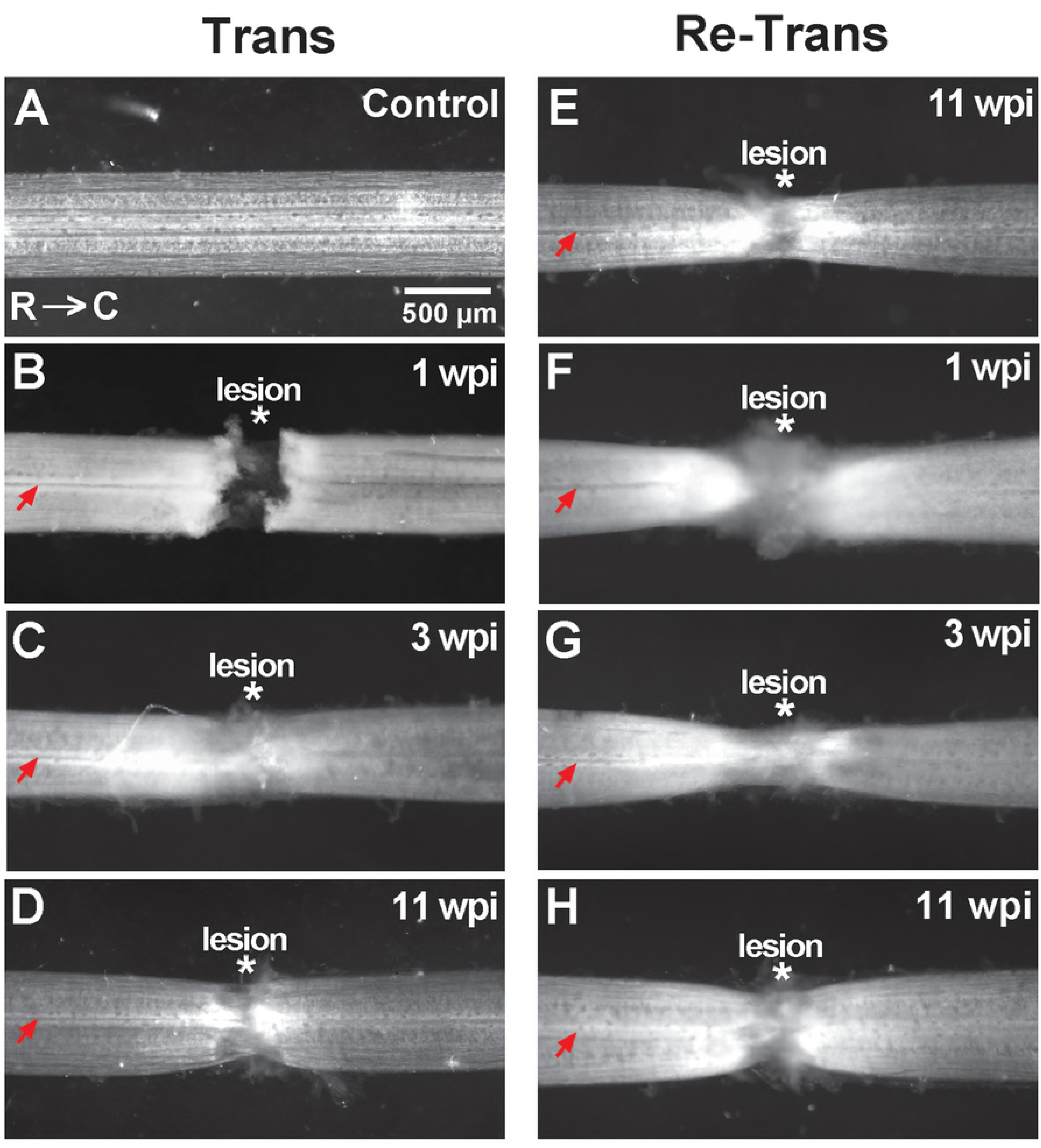
Robust tissue repair after spinal cord transection and re-transection. **(A-D)** Brightfield images showing an uninjured, control lamprey spinal cord (A) and transected spinal cords (B-D) at the indicated time points. At 1wpi the proximal and distal stumps are still separated. But by 3-11 wpi, the spinal cords are reconnected after tissue repair. **(E-H)** Images showing the typical time course of lesion repair within re-transected spinal cords. Asterisks indicate the lesion center, and red arrows indicate the central canal. R=rostral; C=caudal. Scale bar in A applies to B-H.

### Long-distance axon regeneration remains robust after spinal re-transection

Next, we examined the extent of axon regeneration after spinal cord re-transection. To do so, we anterogradely labeled regenerating axons in transected and re-transected lamprey spinal cords using a 10 kDa Alexa Fluor® 488-conjugated fluorescent dextran, as previously described (Lau et al., 2013). Within the uninjured control spinal cord, this procedure preferentially labeled giant RS axons in the ventromedial tract of the spinal cord, as well as medium-caliber fibers in the ventromedial and lateral tracts, the vast majority of which exhibited straight projection patterns (Fig. 3A). In contrast, labeled axons within the transected spinal cord exhibited atypical projection patterns at 11 wpi, including branching and rostral turning, which was especially prevalent in the rostral spinal cord, as previously reported (Fig. 3B, D-E) (Wood and Cohen, 1981; Oliphint et al., 2010). Regenerated axon branches also had smaller diameters (Fig. 3B). Similar morphologies and axonal growth patterns were observed in the re-transected spinal cords at 11 wpi (Fig. 3C, F-G). As a semi-quantitative approach to measuring axon regeneration, we counted the number of labeled axons crossing at 1 mm intervals beyond the lesion center up to 5 mm distal to the lesion (Lau et al., 2013). This analysis showed no significant difference in the number of labeled, regenerated axons in transected and re-transected spinal cords (Fig. 3H) (n = 14 Trans, 7 Re-Trans; ANOVA, p = 0.53). Previous studies in the lamprey model reported that ∼30-70% of descending RS axons regenerate to a position distal to the lesion by 11 wpi (Rovainen, 1976; Yin and Selzer, 1983; Davis and McClellan, 1994; Jacobs et al., 1997). Thus, as a means for comparing with our current findings, we also calculated the percentage of labeled distal/proximal axons and observed ∼50-80% regeneration in the transected spinal cord and no significant differences in the re-transected spinal cords, though the values were slightly higher (Fig. 3I) (n = 14 Trans, 7 Re-Trans; ANOVA, p = 0.94). Thus, axonal re-growth was not impaired by a second spinal transection but remained as robust as after the initial spinal transection.

**Figure 3.**
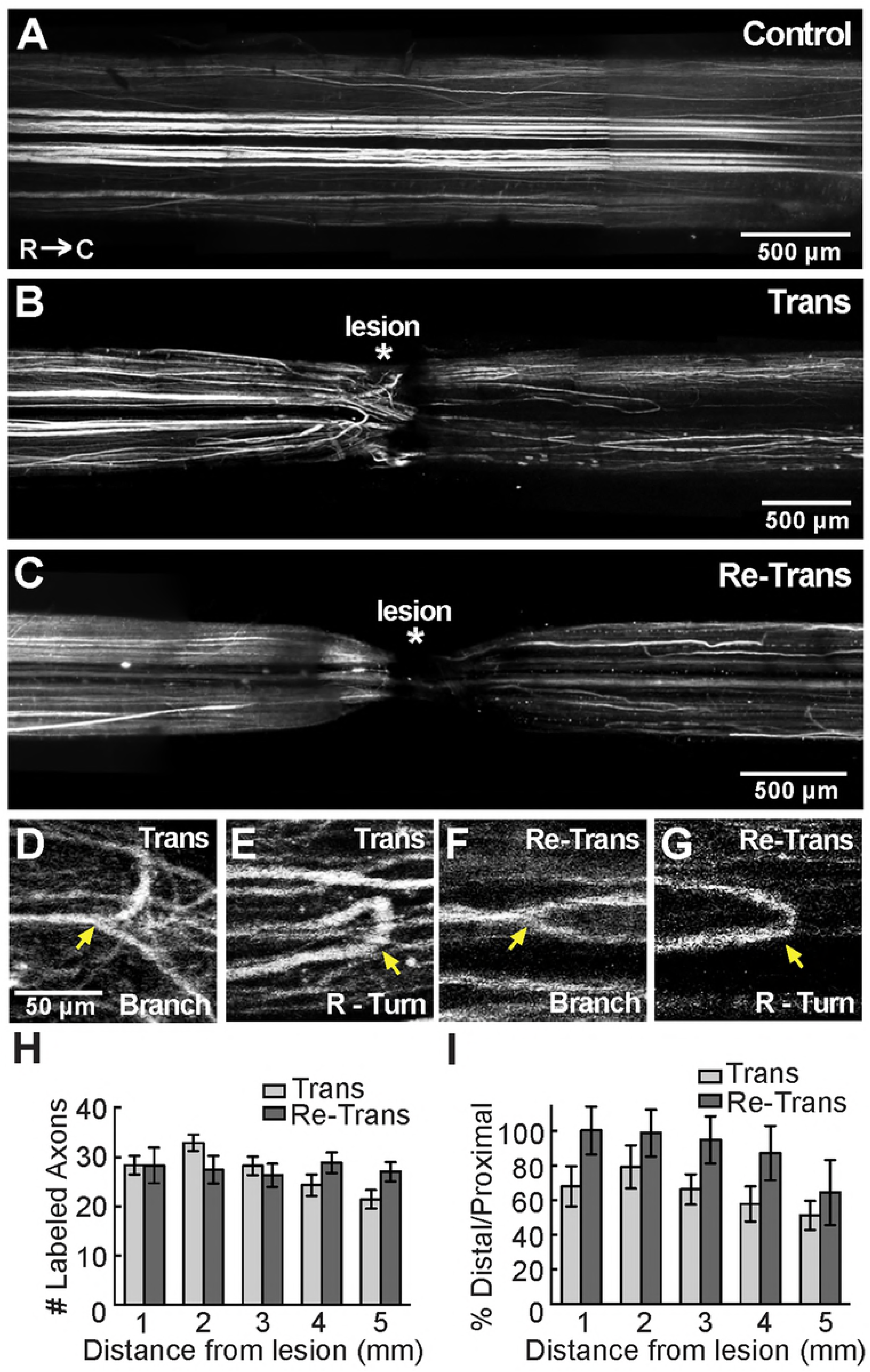
Axon regeneration in the lamprey spinal cord is comparable after repeated transections. **(A)** A montage of confocal z-projections showing bulk labeled axons in the uninjured, control spinal cord. Note the straight axonal projection patterns. (**B-C)** In contrast, regenerated axons in the transected and re-transected spinal cords are sparser at 11 wpi, and they exhibit atypical projection patterns in the medial and lateral tracts. **(D-G)** Higher magnification confocal images showing axonal branching and rostral turning (R-Turn) in transected and re-transected spinal cords (arrows). Scale bar in D applies to E-G. **(H)** There is no significant difference in the number of labeled, regenerated axons in transected and re-transected spinal cords. Bars represent mean ± SEM from n=7-14 lampreys. p = 0.53 by ANOVA. **I.** The percentage of labeled, regenerated axons was also similar. Bars represent mean ± SEM from n=7-14 lampreys. p = 0.13 by ANOVA.

### Axon, synapse, and cytoskeletal distribution is comparable after spinal transection and re-transection

We next examined the distributions of several neuronal and cytoskeletal elements in the transected versus re-transected lamprey spinal cords as another means to assess structural repair. We first immunostained longitudinal cryosections of lamprey spinal cords with a mouse monoclonal antibody against neurofilament-180 (NF-180), which labels large and medium caliber axons in the lamprey spinal cord (Jin et al., 2009; Fogerson et al., 2016). In the control spinal cord, NF-180 staining reveals large and medium-sized RS axons that extend in straight projections through the ventromedial and lateral tracts of the spinal cord (Fig. 4A). DAPI staining labels the central canal, which is narrow and densely packed with nuclei of the ependymal glial cells, as well as motor and intraspinal neurons in the lateral columns (Fig. 4A). At 11 wpi, NF-180 labeling in the transected spinal cord shows regenerating axons that extended through the lesion site and into the distal stump, but with atypical projection patterns (Fig. 4B). In the transected spinal cord, DAPI labeling shows the central canal that is enlarged at and around the lesion site (Fig. 4B). Similar patterns for NF-180 and DAPI staining were observed in the re-transected spinal cord (Fig. 4C). At higher magnification, the altered axonal growth patterns in the transected and re-transected spinal cord can be seen more clearly (Fig. 5A-C). Whereas most axons are straight in the control spinal cord, the axons in the transected and re-transected spinal cords project in winding paths and often cross the spinal cord midline.

**Figure 4.**
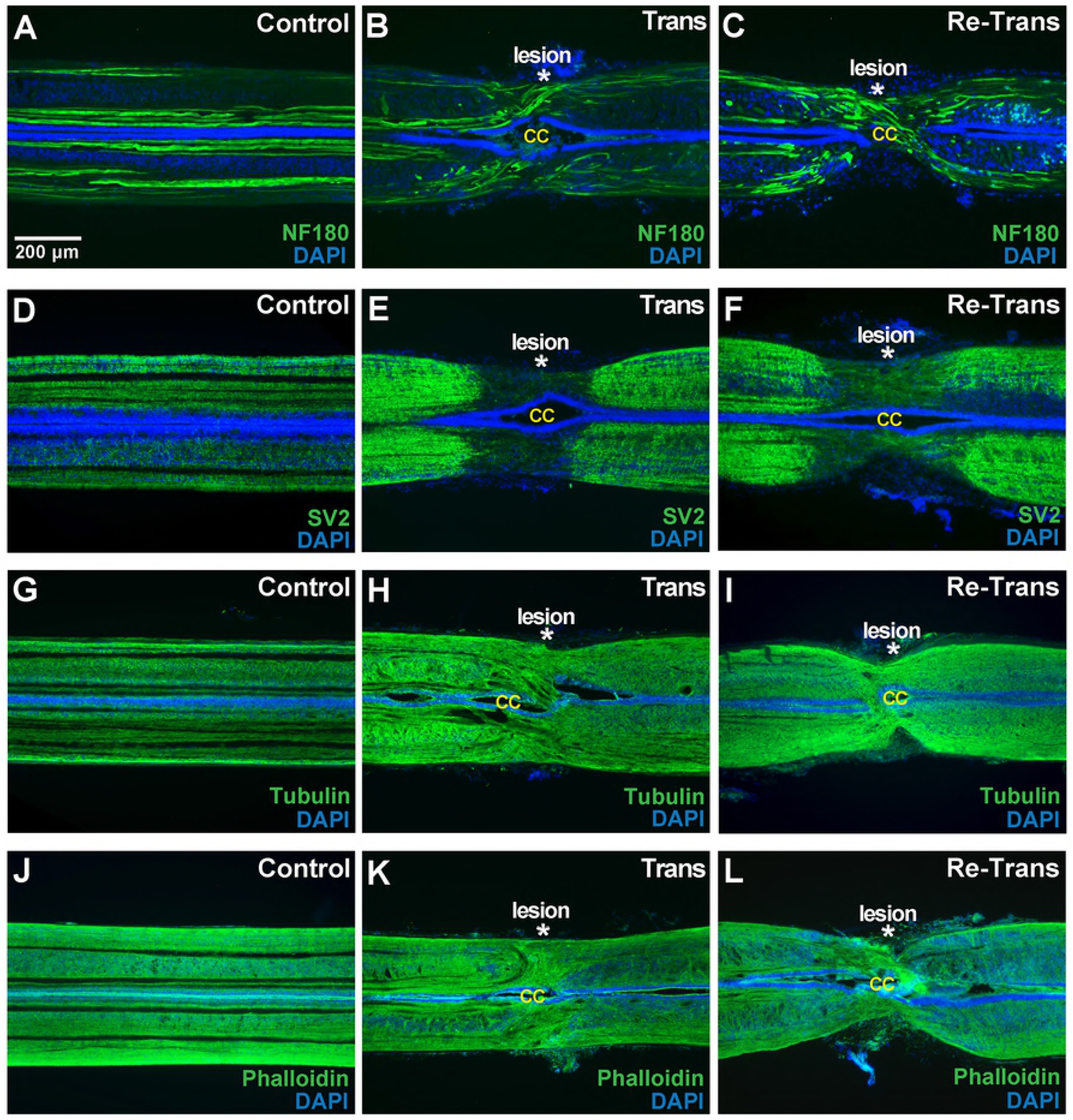
Distributions of axons, synapses, and cytoskeletal elements are similar in lamprey spinal cord after transection and re-transection. **(A-C)** NF-180 antibody labels large and medium caliber axons, which are straight, in the control lamprey spinal cord. DAPI labels nuclei in the central canal (CC), as well as intraspinal neurons in more lateral tracts. In both transected and re-transected spinal cords, NF-180 labels regenerating axons projecting in aberrant patterns. Scale bar in A applies to all panels. Asterisk indicates lesion center. (**D-F)** SV2 is normally distributed in a uniform punctate pattern throughout the control spinal cord. After spinal transection and re-transection, there is a loss of synapses within the lesion site. (**G-L)** Tubulin and phalloidin staining reveals robust microtubule and actin distribution, respectively, throughout the spinal cord in all conditions. There are no obvious differences in labeling between transected and re-transected spinal cords. Rostral is to the left in all images.

**Figure 5.**
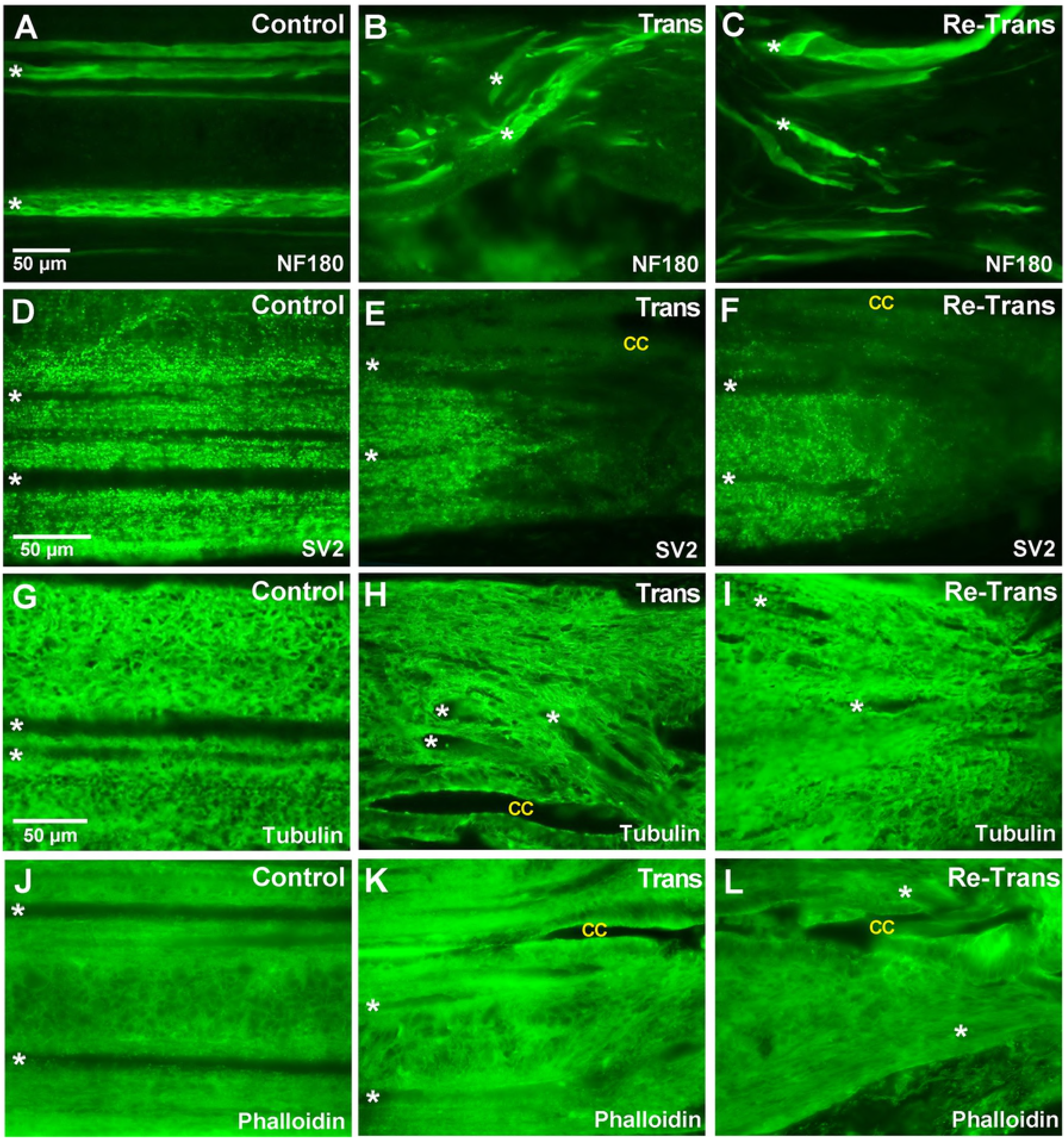
Cellular distribution of axons, synapses, and cytoskeletal elements are comparable after spinal transection and re-transection. **(A-C)** NF-180 staining in large and medium caliber axons in the control, transected and re-transected spinal cords. Note the differences in axonal growth patterns after spinal injury. Asterisks in A-L mark the leftmost position of several identified giant RS axons within each section. Scale bar in A applies to all panels except D and G. **(D-F)** SV2 staining is punctate in all conditions. Note the tapering signal at the rostro-lesion border in the transected and re-transected spinal cords. CC = central canal. **(G-I)** Tubulin staining labels microtubules within neuronal and glial processes throughout the neuropil. There are no major differences after spinal re-transection. **(J-L)** Phalloidin staining labels actin cytoskeleton and appears as a diffuse signal surrounding axons and cell bodies in the spinal cord. Rostral is to the left in all images.

Similarly, spinal cord sections were stained with an antibody against the synaptic vesicle protein SV2, which labels presynaptic vesicle clusters in all vertebrates tested, including lampreys (Buckley and Kelly, 1985; Bloom et al., 2003; Busch and Morgan, 2012; Busch et al., 2014). This allowed us to determine the overall distribution of synapses within the spinal cord. In the uninjured control spinal cord, the SV2 antibody produced fairly uniform, punctate staining throughout the neuropil (Fig. 4D; Fig. 5D). The profiles of large RS axons were visible (dark lines) because their synapses are localized at the periphery along the axolemmal surface. At 11 wpi in the transected spinal cord, the synaptic labeling in the proximal and distal neuropil remained strong, while the density of synapses within the lesion site was markedly decreased, as previously reported (Fig. 4E) (Oliphint et al., 2010). A similar loss of SV2 labeling around the lesion site was also seen within the spinal cords of re-transected animals (Fig. 4F). Higher magnification imaging further revealed SV2 expression at the rostral-lesion border within transected and re-transected spinal cords (Fig. 5E-F).

Additionally, spinal cord sections were immunostained for α-tubulin and phalloidin, which label microtubules and filamentous actin, respectively. The tubulin antibody was a mouse monoclonal that recognized a single band in both rat brain and lamprey CNS lysates at ∼50 kDa, which is the expected molecular weight for tubulin (Suppl. Fig. 1). The α-tubulin and phalloidin labeling were robust and uniform throughout the control spinal cords (Fig. 4G, J), and they revealed web-like structures throughout the neuropil, which are likely to be the intertwining processes of intraspinal neurons and glial cells (Fig. 5G, J). Similar distributions and labeling patterns were observed for both cytoskeletal elements after spinal transection and re-transection, albeit with less organization due to the tissue re-organization in and around the lesion site (Figs. 4H-I, K-L; Fig. 5H-I, K-L). No obvious differences were noted between the tubulin or phalloidin staining in transected versus re-transected cords. Taken together, these data indicate that the distributions of axons, synapses, and cytoskeletal elements within transected lamprey spinal cords are similar after two bouts of regeneration.

### The same subset of giant RS neurons regenerates after spinal transection and re-transection

We next took advantage of the large, identified giant RS neurons as another means to evaluate axon regeneration and neuronal survival after spinal re-transection. The lamprey midbrain and hindbrain contain ∼1200 total RS neurons that descend into the spinal cord and initiate locomotion (Dubuc et al., 2008). Amongst them are 32 giant neurons (100-200 µm in diameter) that reside in stereotypical locations (Fig. 6A) (Rovainen, 1967; Jacobs et al., 1997; Shifman et al., 2008; Busch and Morgan, 2012). These include the mesencephalic cells (M cells: M1-3), isthmic cells (I cells: I1-I5), and bulbar cells (B cells: B1-B6), as well as the Mauthner (Mth) and auxiliary Mauthner (mth’) neurons. These giant RS neurons exhibit distinct intrinsic capacities for surviving and regenerating after axotomy induced by spinal cord transection (Jacobs et al., 1997; Shifman et al., 2008; Busch and Morgan, 2012; Lau et al., 2013; Barreiro-Iglesias, 2015; Fogerson et al., 2016). While some giant RS neurons are “good regenerators” (e.g. M1, I3-I5, B2, B4-B6, mth’), meaning they have a high probability of surviving the injury and regenerating their axons, others are “poor regenerators” (e.g. M2-M3, I1, B1, B3, Mth) with a low probability of survival and regeneration.

**Figure 6.**
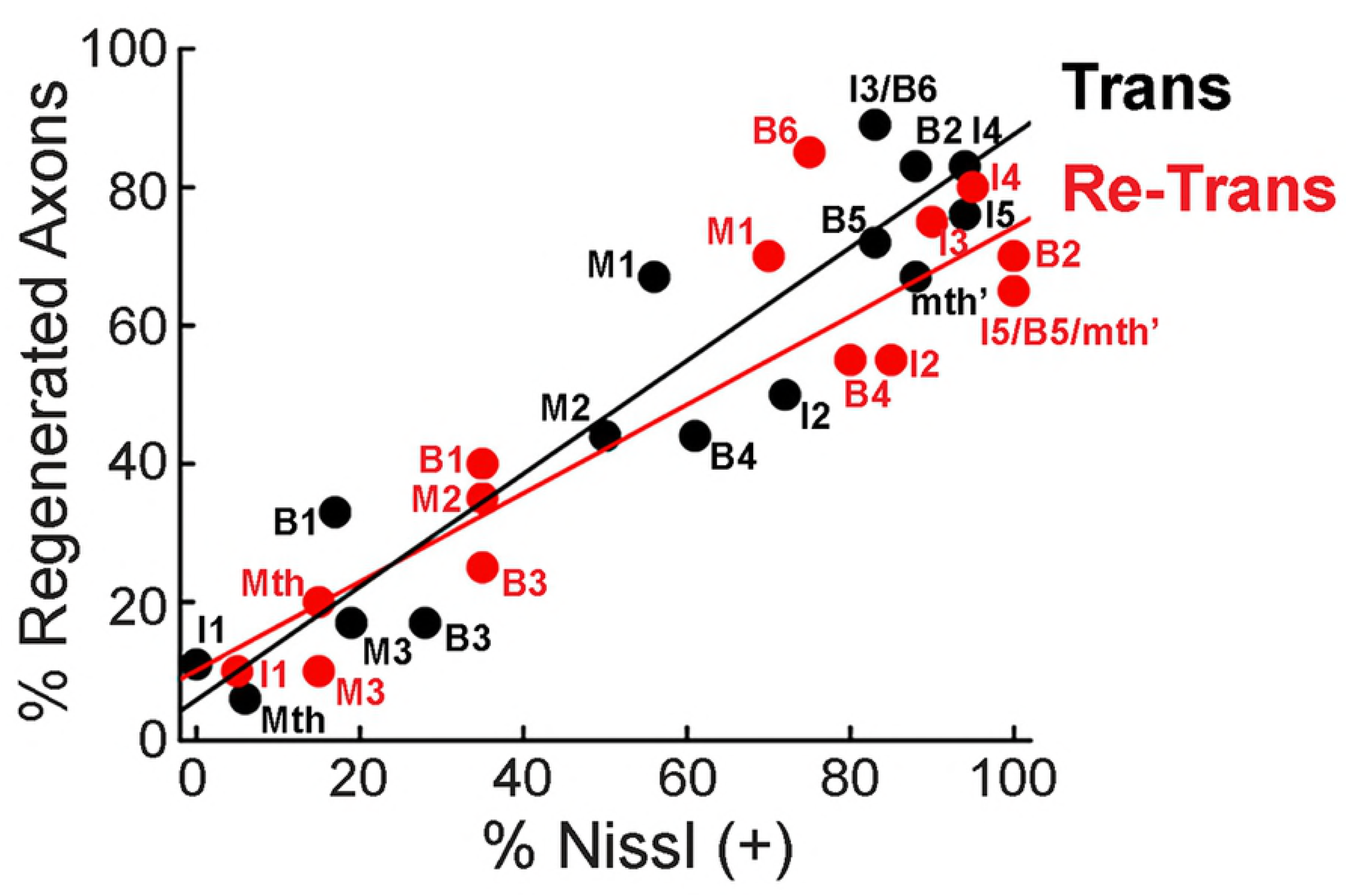
Regeneration of giant RS neurons is indistinguishable after spinal transection and re-transection. **(A)** Diagram showing the giant RS neurons in the lamprey brain. These are the M, I, and B cells, as well as Mauthner neurons. **(B)** Retrograde labeling in a control lamprey reveals all giant RS neurons. **(C)** In contrast, only a subset of giant RS neurons regenerate after spinal transection (white labels), as identified by dye labeling, while others fail to regenerate and are devoid of dye (yellow labels). **(D)** A similar cohort of regenerated neurons is labeled after spinal re-transection. Scale bar in B also applies to C-D. **(E)** Cell-by-cell analysis of giant RS neuron regeneration from a population of n=10 lampreys. There are no obvious differences in regeneration by cell type. **(F)** Similarly, the percentage of all giant RS neurons that regenerated was similar in transected and re-transected lampreys. **(G-H)** Likewise, there were no obvious differences in regeneration of either “good regenerators” or “poor regenerators.” Bars in F and H represent mean ± SEM per brain from n=10 lampreys. n.s. indicates “not significant” by Students t-Test (p>0.05).

We thus wanted to determine whether the giant RS axons retain their normal intrinsic capacities for regeneration after spinal re-transection. To do so, we retrogradely labeled regenerating RS neurons with tetramethylrhodamine applied caudal to the original lesion site (*see Methods*) (Shifman et al., 2008). In the brains of uninjured control animals, all giant RS neurons were labeled using this technique (Fig. 6B). At 11 wpi after a single spinal transection, a select subset of giant RS neurons was labeled, indicating that they had regenerated their axons beyond the lesion site, and these were generally those neurons previously classified as “good regenerators” (Fig. 6C, white labels). In contrast, the remaining giant RS neurons were not labeled, indicating that they did not regenerate their axons, and these were generally those neurons previously classified as “poor regenerators” (Fig. 6C, yellow arrows). A similar pattern of RS neuron labeling was observed in the brains of re-transected lampreys, implicating a similar degree of neuron regeneration (Fig. 6D). Indeed, a cell-by-cell analysis revealed a similar degree of axon regeneration across the giant RS neurons after spinal transection or re-transection (Fig. 6E). At the population level, there was a similar degree of axon regeneration amongst giant RS neurons in transected or re-transected animals (Fig. 6F) (Trans: 53 ± 5% regenerated/brain, n=9 animals; Re-Trans: 52 ± 4% regenerated/brain, n=10 animals; Student’s t-Test, p = 0.88). Likewise, after transection or re-transection, there was a similar degree of regeneration of the “good regenerator” population (i.e. M1, I2-5, B2, B5-6, mth’), which we defined as those giant RS neurons that regenerated >50% of the time (Fig. 6G-H) (*Good Regenerators* - Trans: 75 ± 4% regenerated/brain, n=9 animals, 162 neurons; Re-Trans: 70 ± 3% regenerated/brain, n=10 animals, 180 neurons; Student’s t-Test, p = 0.34). There was a similar amount of regeneration of “poor regenerators” (i.e. M2-3, I1, B1, B3-4, Mth), which we defined as those giant RS neurons that regenerated <50% of the time (*Poor Regenerators* - Trans: 25 ± 6% regenerated/brain, n=9 animals, 108 neurons; Re-Trans: 28 ± 6% regenerated/brain, n=10 animals, 120 neurons; Student’s t-Test, p = 0.71). Thus, the extent of giant RS axon regeneration observed after spinal re-transection was comparable to that after single transections, with the same “good regenerators” exhibiting robust regrowth.

### The same subset of giant RS neurons survive after spinal transection and re-transection

As with axon regeneration, each giant RS neuron exhibits a distinct and reproducible intrinsic capacity for survival or death after axotomy. The “good survivors” (e.g. M1, I2-5, B2, B6, mth’) are those giant RS neurons that typically survive the injury and regenerate their axons, while the “poor survivors” (e.g. M3, I1, B1, B3, Mth) are those that typically undergo delayed death by apoptosis (Shifman et al., 2008; Barreiro-Iglesias and Shifman, 2012; Busch and Morgan, 2012; Fogerson et al., 2016).

As a second measure of giant RS neuron vitality, we evaluated cell survival and death in the same brains that had been previously assayed for axon regeneration (Shifman et al., 2008; Barreiro-Iglesias and Shifman, 2012; Busch and Morgan, 2012). After live imaging to assess the regenerated neurons, as described in Figure 6, we then fixed and histologically stained the brains with Toluidine blue O, which labels Nissl substance. All giant RS neurons within control brains were darkly stained, revealing abundant Nissl substance that is characteristic of healthy neurons (Fig 7A). We refer to these as “Nissl (+)” neurons. In contrast, after a single spinal transection, a subset of neurons (largely the “poor survivors”) had a swollen, chromalytic appearance and lacked Nissl substance, which is indicative of degenerating neurons (Fig. 7B, yellow labels) (Shifman et al., 2008; Busch and Morgan, 2012). We refer to these as “Nissl (-)” neurons. The remaining neurons (largely “good survivors”) retained Nissl substance after spinal transection (Fig. 7B). Following spinal re-transection, a similar Nissl staining pattern was observed amongst the giant RS neurons (Fig. 7C). Across cohorts of transected and re-transected lampreys (n=9-10 animals), the identified giant RS neurons exhibited similar degrees of Nissl (+) staining on a cell-by-cell basis (Fig. 7D, F). There was no significant difference in the percentage of total “Nissl (+)” giant RS neurons per brain after spinal re-transection (Fig. 7E) (Trans: 58 ± 3% Nissl + neurons/brain, n=9 animals, 288 neurons; Re-Trans: 65 ± 3% Nissl (+) neurons/brain, n=10 animals, 320 neurons; Student’s t-Test, p = 0.15). Likewise, there was no significant difference in the percentage of Nissl (+) “good survivors” (i.e. M1, I2-I5, B2, B4-6, mth’), which we defined as those that retained Nissl (+) staining >50% of the time (Fig. 7G) (*Good Survivors:* Trans: 80 ± 4% Nissl (+)/brain, n=9 animals, 180 neurons; Re-Trans: 90 ± 4% Nissl (+)/brain; n=10 animals, 200 neurons; Students t-Test, p = 0.11). Nor were there differences in the percentage of Nissl (+) “poor survivors” (i.e. M2-3, I1, B1, B3, Mth), which we defined as those giant RS neurons that retained Nissl (+) staining <50% of the time (Fig. 7G) (*Poor Survivors:* Trans: 20 ± 7% Nissl (+)/brain, n = 9 animals, 108 neurons; Re-Trans: 23 ± 5% Nissl (+)/brain, n = 10 animals, 120 neurons; Student’s t-Test, p = 0.72). Thus, the probability of survival amongst all giant RS neurons was similar in transected and re-transected lampreys, and the “good survivors” appeared to robustly survive the second injury.

**Figure 7.**
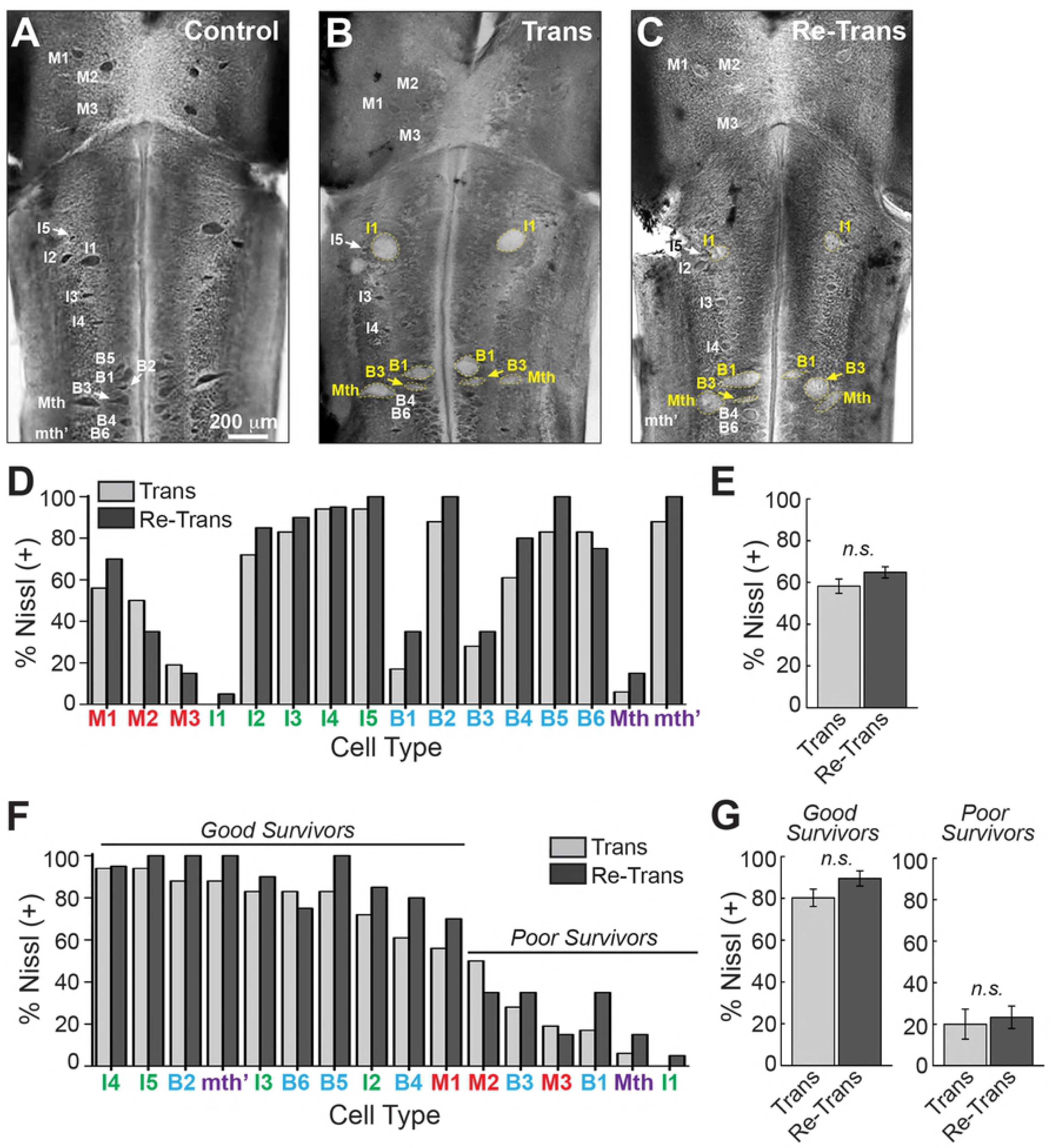
Nissl staining of giant RS neurons is comparable after spinal transection and re-transection. **(A)** A whole mounted control lamprey brain stained with Toluidine blue O, which labels Nissl substance within healthy neurons. All giant RS neurons are labeled. Scale bar in A also applies to B-C. **(B-C)** In contrast, after spinal transection and re-transection only a subset of neurons retains strong Nissl staining (white labels), indicating surviving neurons. Other neurons become chromalytic, swell, and lose their Nissl substance (yellow labels), indicating neurodegeneration. **(D)** Cell-by-cell analysis of Nissl (+) giant RS neurons from n=10 lampreys. There are no obvious differences in Nissl staining by cell type. **(E)** The percentage of total giant RS neurons that was Nissl (+) was similar in transected and re-transected lampreys. **(F-G)** “Good survivor” and “poor survivor” populations of giant RS neurons exhibited similar rates of cell survival, as indicated by Nissl (+) staining. Bars in E and H represent mean ± SEM per brain from n=10 lampreys. n.s. indicates “not significant” by Students T-test (p>0.05).

### The relationship between neurons’ regeneration and survival was unaltered after spinal re-transection

Previous studies have reported a positive linear correlation between neuronal survival (as measured by Nissl staining) and axon regeneration (as measured by retrograde labeling) for the giant RS neurons, such that neurons with a high probability of survival are also likely to regenerate their axons and vice versa (Jacobs et al., 1997; Shifman et al., 2008; Busch and Morgan, 2012; Barreiro-Iglesias, 2015). Using this same approach, we also observed the same positive correlation after a single spinal transection as was previously reported (Fig. 8, black line; R^2^=0.91). The same strong correlation was also observed after spinal re-transection (Fig. 8, red line, R^2^=0.84), which was not significantly different from the former (One-way ANCOVA, p=0.11). Thus, after both spinal cord transection and re-transection, the relationship between neuronal survival and axon regeneration was maintained for each of the giant RS neurons, further corroborating the sustained regenerative potential within lamprey spinal cord.

**Figure 8.**
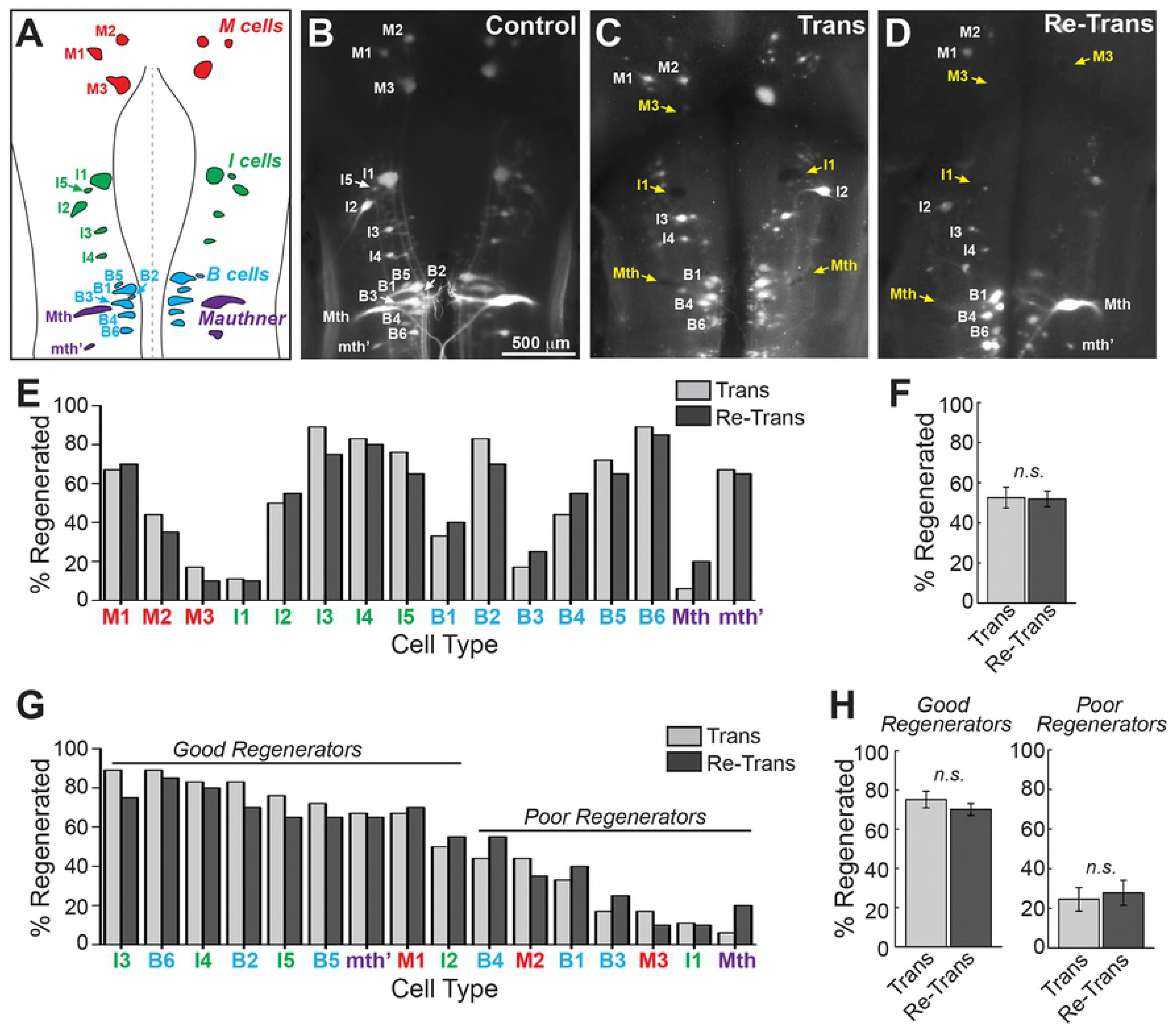
The relationship between cell survival and axon regeneration is similar after spinal re-transection. There is a positive, linear correlation between cell survival [Nissl (+)] and axon regeneration after spinal transection (R^2^ = 0.91), which is also recapitulated after spinal re-transection (R^2^ = 0.84) (ANCOVA p = 0.11).

## Discussion

We report here that the regenerative capacity within the lamprey spinal cord appears to be largely unaltered after two successive transections at the same lesion plane. Behavioral recovery (Fig. 1), tissue repair (Fig. 2), axon regeneration (Fig. 3-4, 6), synapse and cytoskeletal distributions (Fig. 4-5), and cell survival (Fig. 6-7) were nearly identical after recovery from both the first and second spinal transections. Similarly, in axolotls, the area of tail tissue that regenerates after a second amputation is on average the same as after the first amputation, though differences between sexes have been observed (Voss et al., 2013). The Japanese newt (*Cynops pyrrhogaster*) can regenerate a normal lens successively up to 18 times spanning 16 years, demonstrating an unlimited regenerative capacity that is also unaffected by aging (Eguchi et al., 2011). Another striking example is the zebrafish caudal fin, which appears to exhibit unlimited regeneration by regrowing normal fin structures even after 27 amputations (Azevedo et al., 2011). These instances are in stark contrast to repeated limb amputations in amphibians such as axolotls and newts, which exhibit high fidelity regeneration after the first amputation but dramatically decreased regeneration with each successive injury, starting with the second amputation (Dearlove and Dresden, 1976; Bryant et al., 2017). Imperfect limb regeneration in amphibians has also been reported in the fossil record (Frobisch et al., 2014), though naturally the prior status of the limbs cannot be ascertained. Interestingly, in axolotls, performing limb amputations at serially-distal locations resulted in significantly improved regenerative capacity, indicating that the failure of limb regeneration is due to events occurring at the original lesion plane (Bryant et al., 2017). We do not yet understand the full regenerative capacity in the lamprey spinal cord, which would require additional rounds of transection and regeneration. However, the robust, high fidelity regeneration that we observed in the lamprey after two successive spinal cord transections indicates that they have greater regenerative capacity than is observed in some other highly regenerative models.

Functional recovery after spinal cord injury in lampreys and other non-mammalian vertebrates is supported by extensive regeneration of descending axons beyond the lesion scar (Bloom, 2014; Morgan and Shifman, 2014). Previous studies in lampreys reported that 30-70% of descending reticulospinal axons regenerated several millimeters beyond the lesion center by 11 wpi (Yin and Selzer, 1983; Davis and McClellan, 1994; Oliphint et al., 2010; Lau et al., 2013). Results presented here are consistent with this overall level of axon regeneration in the re-transected lamprey spinal cords after the second round of regeneration (Fig. 3I, 6F). The percentages of regenerated axons appears to be slightly higher after bulk anterograde labeling (Fig. 3I), but this is likely due to the possibility of counting multiple branches of the same parent axon that cross at the fiduciary marker.

Remarkably, the cell specificity of axon regeneration amongst the giant RS neurons was also maintained after spinal re-transection (Fig. 6-8). On one hand, it is not surprising that “poor regenerators/survivors” did not regenerate after spinal re-transection, because they had likely undergone delayed degeneration by apoptosis after the first spinal transection, as previously reported (Fig. 6-8) (Shifman et al., 2008; Barreiro-Iglesias and Shifman, 2012; Busch and Morgan, 2012; Barreiro-Iglesias, 2015; Fogerson et al., 2016). However, it is interesting that the extent of axon regeneration and cell survival of the remaining RS neurons (e.g. M1, I2-I5, B2, B5-B6, mth’) was nearly the same after both spinal transection and re-transection (Fig. 6-8). It is unlikely that the giant RS neurons were replaced by newly-born neurons after transection or re-transection, as neurogenesis appears to be fairly limited in the brain after spinal injury and also restricted to the ependymal zone (Zhang et al., 2014). It is thus likely that many of the “good regenerators/survivors” underwent two rounds of regeneration during the 22-week experiment, suggesting that the intrinsic regenerative capacity of individual giant RS neurons was also largely unaffected by spinal re-transection. Determining this unequivocally would require long-term dynamic imaging in the lamprey nervous system, which is not yet practical in our system.

While lampreys recover normal swimming behaviors after spinal cord transection and re-transection, it must be acknowledged that functional recovery is the result of substantial plasticity throughout the central nervous system. That is, the regenerated spinal cord does not return to the original status of an uninjured spinal cord but rather forms new functional circuitry with compensatory network properties (Parker, 2017). This is clearly illustrated by the facts that only a subset of descending axons regenerate in the transected and re-transected spinal cord (Fig. 3, 6), and those that regenerate terminate early, exhibit atypical projection patterns, and produce few synapses (Fig. 3, 4A-F) (Wood and Cohen, 1981; Yin et al., 1981; Yin and Selzer, 1983; Oliphint et al., 2010). Yet, the excitatory postsynaptic potentials, a measure of synaptic strength, can be as strong or stronger than those in the uninjured spinal cord (Mackler and Selzer, 1985, 1987). In addition, using electrophysiological methods, compensatory plasticity has also been documented at regenerated lamprey spinal synapses, as are changes in the intrinsic properties of regenerated axons, which together could boost the synaptic output of regenerated synapses (Cooke and Parker, 2009; Parker, 2017). Given the remarkable consistency of axon, synapse and cytoskeleton distributions (Fig. 4-5); axon regeneration (Fig. 3, 6); and cell survival (Fig. 7) in the transected and re-transected spinal cords, it is likely that the second bout of regeneration induces similar types of neural plasticity, though this remains to be fully explored. The one obvious difference we observed was at 1wpi where the re-transected spinal cord appeared to have accelerated the formation of scar tissue (Fig. 2B, F).

Going forward, it will be important to further investigate and compare the cellular and molecular mechanisms of tissue repair and regeneration after spinal transection and re-transection. Doing so would allow us to identify how regenerative capacity remains as robust after additional injuries. RNA-Seq analysis, such as that which was recently performed on singly transected spinal cords (Herman et al., 2018), may therefore be useful as an unbiased means for beginning to identify these mechanisms in the re-transected spinal cords. Doing so will permit a greater understanding of the molecular requirements that are driving successful regeneration of the vertebrate central nervous system and may provide insights into the limitations that occur in non-regenerative conditions.

## Acknowledgements

The authors would like to thank Dr. Cristina Roman-Vendrell and Louie Kerr, Director of the Central Microscopy Facility at the MBL, for technical support. We also thank Dr. Juan Diaz-Quiroz for helpful comments on the manuscript. EG was supported in part by an NSF REU Award (#1659604: Biological Discovery in Woods Hole at the Marine Biological Laboratory).

## Supporting Information

**Figure S1.Characterization of α-tubulin antibody.** Western blot using a mouse monoclonal α-tubulin antibody (Sigma; clone DM1A) revealed a single band in both rat brain and lamprey CNS lysates at ∼50 kDa, which is the expected molecular weight for α-tubulin.

## References

Azevedo AS, Grotek B, Jacinto A, Weidinger G, Saude L (2011) The regenerative capacity of the zebrafish caudal fin is not affected by repeated amputations. PloS one 6:e22820.

Barreiro-Iglesias A (2015) “Bad regenerators” die after spinal cord injury: insights from lampreys. Neural Regen Res 10:25–27.

Barreiro-Iglesias A, Shifman MI (2012) Use of fluorochrome-labeled inhibitors of caspases to detect neuronal apoptosis in the whole-mounted lamprey brain after spinal cord injury. Enzyme research 2012:835731.

Bely AE (2014) Early events in annelid regeneration: a cellular perspective. Integr Comp Biol 54:688–699.

Bloom O (2014) Non-mammalian model systems for studying neuro-immune interactions after spinal cord injury. Exp Neurol 258:130–140.

Bloom O, Evergren E, Tomilin N, Kjaerulff O, Low P, Brodin L, Pieribone VA, Greengard P, Shupliakov O (2003) Colocalization of synapsin and actin during synaptic vesicle recycling. J Cell Biol 161:737–747.

Bryant DM, Sousounis K, Farkas JE, Bryant S, Thao N, Guzikowski AR, Monaghan JR, Levin M, Whited JL (2017) Repeated removal of developing limb buds permanently reduces appendage size in the highly-regenerative axolotl. Developmental biology 424:1–9.

Buckley K, Kelly RB (1985) Identification of a transmembrane glycoprotein specific for secretory vesicles of neural and endocrine cells. J Cell Biol 100:1284–1294.

Busch DJ, Morgan JR (2012) Synuclein accumulation is associated with cell-specific neuronal death after spinal cord injury. J Comp Neurol 520:1751–1771.

Busch DJ, Oliphint PA, Walsh RB, Banks SM, Woods WS, George JM, Morgan JR (2014) Acute increase of alpha-synuclein inhibits synaptic vesicle recycling evoked during intense stimulation. Mol Biol Cell 25:3926–3941.

Cohen AH, Mackler SA, Selzer ME (1986) Functional regeneration following spinal transection demonstrated in the isolated spinal cord of the larval sea lamprey. Proc Natl Acad Sci U S A 83:2763–2766.

Cooke RM, Parker D (2009) Locomotor recovery after spinal cord lesions in the lamprey is associated with functional and ultrastructural changes below lesion sites. J Neurotrauma 26:597–612.

David S, Aguayo AJ (1981) Axonal elongation into peripheral nervous system “bridges” after central nervous system injury in adult rats. Science 214:931–933.

Davis GR, Jr., McClellan AD (1994) Long distance axonal regeneration of identified lamprey reticulospinal neurons. Exp Neurol 127:94–105.

Davis GR, Jr., Troxel MT, Kohler VJ, Grossmann EM, McClellan AD (1993) Time course of locomotor recovery and functional regeneration in spinal-transected lamprey: kinematics and electromyography. Exp Brain Res 97:83–95.

Dearlove GE, Dresden MH (1976) Regenerative abnormalities in Notophthalmus viridescens induced by repeated amputations. J Exp Zool 196:251–262.

Diaz Quiroz JF, Echeverri K (2013) Spinal cord regeneration: where fish, frogs and salamanders lead the way, can we follow? Biochem J 451:353–364.

Dolan CP, Dawson LA, Muneoka K (2018) Digit Tip Regeneration: Merging Regeneration Biology with Regenerative Medicine. Stem Cells Transl Med 7:262–270.

Dubuc R, Brocard F, Antri M, Fenelon K, Gariepy JF, Smetana R, Menard A, Le Ray D, Viana Di Prisco G, Pearlstein E, Sirota MG, Derjean D, St-Pierre M, Zielinski B, Auclair F, Veilleux D (2008) Initiation of locomotion in lampreys. Brain Res Rev 57:172–182.

Echeverri K, Tanaka EM (2002) Ectoderm to mesoderm lineage switching during axolotl tail regeneration. Science 298:1993–1996.

Eguchi G, Eguchi Y, Nakamura K, Yadav MC, Millan JL, Tsonis PA (2011) Regenerative capacity in newts is not altered by repeated regeneration and ageing. Nature communications 2:384.

Erickson JR, Gearhart MD, Honson DD, Reid TA, Gardner MK, Moriarity BS, Echeverri K (2016) A novel role for SALL4 during scar-free wound healing in axolotl. NPJ Regen Med 1.

Fawcett JW, Gaze RM (1981) The organization of regenerating axons in the Xenopus optic nerve. Brain Res 229:487–490.

Fogerson SM, van Brummen AJ, Busch DJ, Allen SR, Roychaudhuri R, Banks SM, Klarner FG, Schrader T, Bitan G, Morgan JR (2016) Reducing synuclein accumulation improves neuronal survival after spinal cord injury. Exp Neurol 278:105–115.

Frobisch NB, Bickelmann C, Witzmann F (2014) Early evolution of limb regeneration in tetrapods: evidence from a 300-million-year-old amphibian. Proc Biol Sci 281:20141550.

Gemberling M, Bailey TJ, Hyde DR, Poss KD (2013) The zebrafish as a model for complex tissue regeneration. Trends Genet 29:611–620.

Gibbs KM, Chittur SV, Szaro BG (2011) Metamorphosis and the regenerative capacity of spinal cord axons in Xenopus laevis. Eur J Neurosci 33:9–25.

Goldshmit Y, Sztal TE, Jusuf PR, Hall TE, Nguyen-Chi M, Currie PD (2012) Fgf-dependent glial cell bridges facilitate spinal cord regeneration in zebrafish. J Neurosci 32:7477–7492.

Gorsuch RA, Hyde DR (2014) Regulation of Muller glial dependent neuronal regeneration in the damaged adult zebrafish retina. Exp Eye Res 123:131–140.

Herman PE, Papatheodorou A, Bryant SA, Waterbury CKM, Herdy JR, Arcese AA, Buxbaum JD, Smith JJ, Morgan JR, Bloom O (2018) Highly conserved molecular pathways, including Wnt signaling, promote functional recovery from spinal cord injury in lampreys. Sci Rep 8:742.

Jacobs AJ, Swain GP, Snedeker JA, Pijak DS, Gladstone LJ, Selzer ME (1997) Recovery of neurofilament expression selectively in regenerating reticulospinal neurons. J Neurosci 17:5206–5220.

Jin LQ, Zhang G, Jamison C, Jr., Takano H, Haydon PG, Selzer ME (2009) Axon regeneration in the absence of growth cones: acceleration by cyclic AMP. J Comp Neurol 515:295–312.

Kang H, Lichtman JW (2013) Motor axon regeneration and muscle reinnervation in young adult and aged animals. J Neurosci 33:19480–19491.

Lau BY, Fogerson SM, Walsh RB, Morgan JR (2013) Cyclic AMP promotes axon regeneration, lesion repair and neuronal survival in lampreys after spinal cord injury. Exp Neurol 250:31–42.

Lau BY, Foldes AE, Alieva NO, Oliphint PA, Busch DJ, Morgan JR (2011) Increased synapsin expression and neurite sprouting in lamprey brain after spinal cord injury. Exp Neurol 228:283–293.

Lurie DI, Pijak DS, Selzer ME (1994) Structure of reticulospinal axon growth cones and their cellular environment during regeneration in the lamprey spinal cord. J Comp Neurol 344:559–580.

Mackler SA, Selzer ME (1985) Regeneration of functional synapses between individual recognizable neurons in the lamprey spinal cord. Science 229:774–776.

Mackler SA, Selzer ME (1987) Specificity of synaptic regeneration in the spinal cord of the larval sea lamprey. J Physiol 388:183–198.

McCusker C, Gardiner DM (2011) The axolotl model for regeneration and aging research: a mini-review. Gerontology 57:565–571.

Monaghan JR, Walker JA, Page RB, Putta S, Beachy CK, Voss SR (2007) Early gene expression during natural spinal cord regeneration in the salamander Ambystoma mexicanum. J Neurochem 101:27–40.

Morgan J, Shifman MI (2014) Non-mammalian models of nerve regeneration. Cambridge, UK: Cambridge University Press.

Morgan TH (1898) Experimental studies of the regeneration of Planaria maculata. Arch Entw Mech Org 7:364–397.

Muneoka K, Bryant SV (1982) Evidence that patterning mechanisms in developing and regenerating limbs are the same. Nature 298:369–371.

Newmark PA, Sanchez Alvarado A (2002) Not your father’s planarian: a classic model enters the era of functional genomics. Nat Rev Genet 3:210–219.

Oliphint PA, Alieva N, Foldes AE, Tytell ED, Lau BY, Pariseau JS, Cohen AH, Morgan JR (2010) Regenerated synapses in lamprey spinal cord are sparse and small even after functional recovery from injury. J Comp Neurol 518:2854–2872.

Parker D (2017) The Lesioned Spinal Cord Is a “New” Spinal Cord: Evidence from Functional Changes after Spinal Injury in Lamprey. Front Neural Circuits 11:84.

Reddien PW, Sanchez Alvarado A (2004) Fundamentals of planarian regeneration. Annu Rev Cell Dev Biol 20:725–757.

Reginelli AD, Wang YQ, Sassoon D, Muneoka K (1995) Digit tip regeneration correlates with regions of Msx1 (Hox 7) expression in fetal and newborn mice. Development 121:1065– 1076.

Rovainen CM (1967) Physiological and anatomical studies on large neurons of central nervous system of the sea lamprey (Petromyzon marinus). I. Muller and Mauthner cells. J Neurophysiol 30:1000–1023.

Rovainen CM (1976) Regeneration of Muller and Mauthner axons after spinal transection in larval lampreys. J Comp Neurol 168:545–554.

Seifert AW, Monaghan JR, Voss SR, Maden M (2012) Skin regeneration in adult axolotls: a blueprint for scar-free healing in vertebrates. PloS one 7:e32875.

Selzer ME (1978) Mechanisms of functional recovery and regeneration after spinal cord transection in larval sea lamprey. J Physiol 277:395–408.

Shifman MI, Zhang G, Selzer ME (2008) Delayed death of identified reticulospinal neurons after spinal cord injury in lampreys. J Comp Neurol 510:269–282.

Slack JM, Lin G, Chen Y (2008) The Xenopus tadpole: a new model for regeneration research. Cell Mol Life Sci 65:54–63.

Son YJ, Thompson WJ (1995) Schwann cell processes guide regeneration of peripheral axons. Neuron 14:125–132.

Sperry RW (1947) Nature of functional recovery following regeneration of the oculomotor nerve in amphibians. Anat Rec 97:293–316.

Tanaka EM, Ferretti P (2009) Considering the evolution of regeneration in the central nervous system. Nat Rev Neurosci 10:713–723.

Tanaka EM, Reddien PW (2011) The cellular basis for animal regeneration. Dev Cell 21:172–185.

Uygur A, Lee RT (2016) Mechanisms of Cardiac Regeneration. Dev Cell 36:362–374.

Vergara MN, Del Rio-Tsonis K (2009) Retinal regeneration in the Xenopus laevis tadpole: a new model system. Mol Vis 15:1000–1013.

Voss GJ, Kump DK, Walker JA, Voss SR (2013) Variation in salamander tail regeneration is associated with genetic factors that determine tail morphology. PloS one 8:e67274.

Voss SR, Epperlein HH, Tanaka EM (2009) Ambystoma mexicanum, the axolotl: a versatile amphibian model for regeneration, development, and evolution studies. Cold Spring Harb Protoc 2009:pdb emo128.

Wang S, Miller SR, Ober EA, Sadler KC (2017) Making It New Again: Insight Into Liver Development, Regeneration, and Disease From Zebrafish Research. Curr Top Dev Biol 124:161–195.

Williams RR, Venkatesh I, Pearse DD, Udvadia AJ, Bunge MB (2015) MASH1/Ascl1a leads to GAP43 expression and axon regeneration in the adult CNS. PloS one 10:e0118918.

Wood MR, Cohen MJ (1981) Synaptic regeneration and glial reactions in the transected spinal cord of the lamprey. J Neurocytol 10:57–79.

Yin HS, Selzer ME (1983) Axonal regeneration in lamprey spinal cord. J Neurosci 3:1135–1144.

Yin HS, Wellerstein KK, Selzer ME (1981) Effects of axotomy on lamprey spinal neurons. Exp Neurol 73:750–761.

Zhang G, Vidal Pizarro I, Swain GP, Kang SH, Selzer ME (2014) Neurogenesis in the lamprey central nervous system following spinal cord transection. J Comp Neurol 522:1316–1332.

Zukor KA, Kent DT, Odelberg SJ (2011) Meningeal cells and glia establish a permissive environment for axon regeneration after spinal cord injury in newts. Neural Dev 6:1.

